# Leveraging breeding values obtained from random regression models for genetic inference of longitudinal traits

**DOI:** 10.1101/435685

**Authors:** Malachy Campbell, Mehdi Momen, Harkamal Walia, Gota Morota

## Abstract

Understanding the genetic basis of dynamic plant phenotypes has largely been limited due to lack of space and labor resources needed to record dynamic traits, often destructively, for a large number of genotypes. However, the recent advent of image-based phenotyping platforms has provided the plant science community with an effective means to non-destructively evaluate morphological, developmental, and physiological processes at regular, frequent intervals for a large number of plants throughout development. The statistical frameworks typically used for genetic analyses (e.g. genome-wide association mapping, linkage mapping, and genomic prediction) in plant breeding and genetics are not particularly amenable for repeated measurements. Random regression (RR) models are routinely used in animal breeding for the genetic analysis of longitudinal traits, and provide a robust framework for modeling traits trajectories and performing genetic analysis simultaneously. We recently used a RR approach for genomic prediction of shoot growth trajectories in rice using 33,674 SNPs. In this study, we have extended this approach for genetic inference by leveraging genomic breeding values derived from RR models for rice shoot growth during early vegetative development. This approach provides improvements over a conventional single time point analyses for discovering loci associated with shoot growth trajectories. The RR approach uncovers persistent, as well as time-specific, transient quantitative trait loci. This methodology can be widely applied to understand the genetic architecture of other complex polygenic traits with repeated measurements.

Core Ideas:
- Random regression models are an appealing framework for GWAS of longitudinal traits
- This approach provides improvements over a conventional single time point analyses for GWAS
- We identify QTL with transient and persistent effects on shoot growth in rice

## 1 Introduction

A plant’s phenotype at any given time is the manifestation of numerous biological processes that have occurred prior to the capture of the phenotype. In most genetic mapping studies, plants are phenotyped at one or few discrete time points. While this may be sufficient for end point traits, such as yield or grain quality, other agronomically important traits such as plant height or vigor are not static and vary continuously throughout development. Given the dynamic nature of these traits, it is likely that some genes will have a time-dependent contribution to the phenotype. Approaches that consider such infinite-dimensional traits as static, fail to fully capture the dynamic processes that have led to the phenotype and may not uncover the contributions of time-specific loci.

Recording phenotypic measurements across development in genetic mapping populations is typically limited due to high space and labor demands to record a trait, often destructively, for a large number of genotypes. However, with the advent of image-based phenotyping platforms, researchers can now capture morphological, developmental, and physiological processes non-destructively with higher temporal resolution for a large number of plants (Fraas and Lüthen, 2015; Simko et al., 2016; Shakoor et al., 2017; Tardieu et al., 2017; Araus et al., 2018). Moreover, the growth of the unmanned aerial vehicle industry in recent years has provided many low-cost hardware options that can be outfitted with cameras, facilitating the collection of temporal phenotypes in field settings (Yang et al., 2017). While the use of these platforms is becoming more routine in plant genetics, the statistical frameworks typically used for genetic analyses (e.g. genome-wide association mapping, linkage mapping, and genomic prediction) in plant breeding and genetics are not amenable for longitudinal traits.

Several studies in recent years have sought to elucidate the genetic basis of longitudinal traits through genome-wide association studies (GWAS) or linkage mapping. For instance Moore et al. (2013) and Würschum et al. (2014) utilized linkage mapping at discrete time points to identify time-specific quantitative trait loci (QTL) associated with root gravitropism and plant height, respectively. While these approaches may be effective, by considering the phenotype at only a single time point they do not leverage the covariance among time points and may have reduced statistical power compared to approaches that consider the entire trait trajectory in regression modeling. Several studies have leveraged a “two-step” approach for functional association mapping (Bac-Molenaar et al., 2015; Campbell et al., 2017). In the two-step approach, a function is fit to phenotypic records for each genotype that summarizes the trait trajectories using a few parameters. These parameters are then used as derived phenotypes in subsequent GWAS analyses. However, with these “two-step” approaches information is lost between the curve fitting and genetic analysis steps. The residuals from the first curve-fitting step likely contain important information regarding persistent environmental effects that are not considered in subsequent genetic analysis. We hypothesize that an approach that unifies the curve fitting and genetic analysis into a single framework is likely to be better than the single time point or a “two-step” longitudinal approach.

Random regression (RR) models provide a robust framework for modeling trait trajectories and performing genetic analysis simultaneously (Schaeffer, 1994; Huisman et al., 2002; Schaeffer, 2004; Sun et al., 2017). Covariance functions, such as spline or polynomial functions, are used to model trait trajectories for each line and sufficiently capture the covariance across time points while estimating fewer parameters (Kirkpatrick et al., 1990; Meyer, 1998; White et al., 1999; Strabel and Misztal, 1999; Pool et al., 2000; Huisman et al., 2002; Schaeffer, 2004; Misztal, 2006; Sun et al., 2017). In a recent study Sun et al. (2017) utilized a RR approach with cubic splines in wheat to obtain best linear unbiased predictions of secondary traits derived from high-throughput hyperspectral and thermal imaging. Regression coefficients are treated as random effects, and therefore allow values to vary between individuals. Genomic estimated breeding values (GEBVs) for regression coefficients are obtained using a mixed model, and using simple algebra, GEBVs can be obtained for any time throughout the continuous trait trajectory (Mrode, 2014)

GEBVs represent the summation of all additive genetic effects across the genome for a given individual. Goddard (2009) showed that GEBVs predicted using genomic relationships (e.g. genomic best linear unbiased prediction (gBLUP)) are equivalent to those predicted from regression on markers. Given this equivalence, marker effects can be easily calculated from GEBVs, thus genetic inference (e.g. GWAS) can be performed. While this approach is different compared to conventional single marker regression GWAS (SMR-GWAS) approaches, it offers several advantages. First, 100,000s of statistical tests are typically run for SMR-GWAS, and as a result, a stringent *p*-value threshold must be used to limit false discoveries (Hayes, 2013). Thus, loci recovered using SMR-GWAS approaches typically account for only a fraction of the total genetic variance for a trait (Yang et al., 2010). Whole-genome BLUP approaches (i.e. SNP-BLUP or gBLUP) assume an infinitesimal model in which all loci have some, albeit small, contribution to the phenotype (Hayes, 2013). Thus, by considering all markers simultaneously small-effect QTL are recovered and more genetic variation can be captured compared to SMR-GWAS (Yang et al., 2010). BLUP approaches shrink marker effects towards zero, and thus may not be appropriate for simple traits that are regulated by few loci with large effects. However, for complex polygenic traits these assumptions are reasonable and should yield biologically meaningful results. In the case of RR, GEBVs can be calculated at each time point and can be leveraged to examine the contribution of loci across a trait trajectory or the time axis.

In a recent study, we used a RR approach for genomic prediction of shoot growth trajectories in rice (Campbell et al., 2018). The utilization of longitudinal phenotypes with RR captured greater genetic variation compared to single time point approach, and significantly improved prediction accuracy. In the current study, we have leveraged GEBVs derived from RR models to examine the genetic architecture of shoot growth through a 20-day period during early vegetative development. We show that this approach can be used for genetic inference of shoot growth trajectories and uncovers persistent, as well as time-specific QTL. Furthermore, we show that the RR approach uncovers considerably more associations compared to a conventional single time point analysis.

## 2 Materials and Methods

### 2.1 High-throughput phenotyping

Phenotypic data was collected for 357 diverse rice accessions from the Rice Diversity Panel 1 (RDP1) (Zhao et al., 2011). The plant materials, experimental design, and image processing are described in detail in Campbell et al. (2018). Briefly, 378 lines were phenotyped at the Plant Accelerator, Australian Plant Phenomics Facility, at the University of Adelaide, SA, Australia from February to April 2016. In this period, three experiments were conducted where experiment consisted of a partially replicated design with 54 randomly selected lines having two replicates in each experiment. The plants were grown on greenhouse benches for 10 days after transplanting (DAT) and were loaded on the imaging system and watered to 90% field capacity at 11 DAT.

Briefly, 378 lines were phenotyped at the Plant Accelerator, Australian Plant Phenomics Facility, at the University of Adelaide, SA, Australia from February to April 2016. In this period, three experiments were conducted where experiment consisted of a partially replicated design with 54 randomly selected lines having two replicates in each experiment. The plants were grown on greenhouse benches for 10 days after transplanting (DAT) and were loaded on the imaging system and watered to 90% field capacity at 11 DAT.

The plants were imaged daily from 13 to 33 DAT using a visible (red–green–blue camera; Basler Pilot piA2400–12 gc, Ahrensburg, Germany) from two side-view angles separated by 90° and a single top view. The LemnaGrid software was used to extract “plant pixels” from the RGB images using a color classification strategy, and noise (i.e. small areas of non-plant pixels) in the image were removed using a series of erosion and dilation steps. Projected shoot area (PSA) was calculated as the sum of the plant areas projected in two dimensional space from each of the three RGB images, and was used as a measure of shoot biomass. Previous studies have shown a high correlation between PSA and conventional destructive measures of shoot biomas (Golzarian et al., 2011; Campbell et al., 2015; Neilson et al., 2015; Knecht et al., 2016). A depiction of PSA collected from RGB images is provided as Figure S1. Outlier plants at each time point were detected at each time point using the 1.5(IQR) rule. Briefly, the distribution of PSA at each day was split into quartiles and the interquartile range (IQR) was calculated as the difference between the third and first quartiles. Points that were either less than Q1 – 1.5 (IQR) or Q3 + 1.5(IQR) were considered as outliers. Outliers were plotted and those that exhibited abnormal growth patterns were removed. A total of 2,604 plants remained for downstream analyses.

### 2.2 Predicting genomic breeding values

#### 2.2.1 Random regression

Trajectories for PSA across the 20-time points was modeled using a RR model with Legendre polynomials. The model is the same that was used for genomic prediction in Campbell et al. (2018). The model is described below using the notation of Mrode (2014)

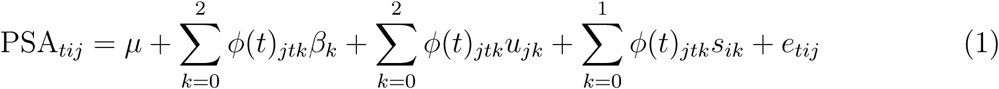

PSA_*tij*_ is PSA on day *t* for line *j* within experiment *i*. _*k*_ is the fixed second-order Legendre polynomial to model the mean PSA trajectory for all lines, *u*_*jk*_ and *s*_*ik*_ are the *k*^*th*^ random regression coefficients for additive genetic effect and random experiment effects, and *e*_*tij*_ is the random residual. The order of *β* was selected based on visual inspection of the PSA over the 20 days. The random additive genetic effects (*u*) are modeled using a second-order Legendre polynomial, and the experiment effects (*s*) are modeled using a first-order Legendre polynomial.

In matrix notation, the model is

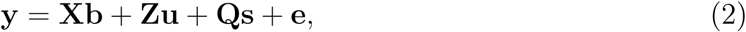

**y** is a vector of PSA over the 20 days and is of order *n*, where *n* is the number of observations. **X** is an *n* × *k*_*f*_ covariable matrix where the number of columns is equal to the order of Legendre polynomial used to model fixed effects (*k*_*f*_). The matrices **Z** and **Q** are covariable matrices for the random additive genetic and random experimental effects, respectively. The number of rows for **Z** is *n* and the number of columns corresponds to the order of Legendre polynomial times the number of lines used to fit the additive genetic effect (*q* * *k*_*g*_ = 357 * 3 = 1, 071). The dimension of **Q** is *n* × *e* * *k*_*s*_ where *k*_*s*_ is the order of Legendre polynomial used to fit the permanent environmental effects and *e* is the number of experiments. We assume **u** ∼ *N* (0, **G** ⊗ **Ω**), **s** ∼ *N* (0, **I** ⊗ **P**), and **e** ∼ *N* (0, **I** ⊗ **D**). Here, **Ω** and **P** are the covariance matrices for the RR coefficients for the additive genetic and permanent environmental effects, and **D** is a diagonal matrix that allows for heterogeneous variances over the 20-time points.

A genomic relationship matrix (**G**) was calculated using VanRaden (2008).

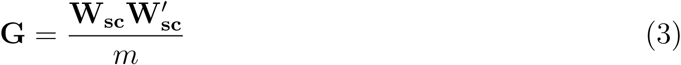

**W_sc_** is a centered and scaled *q* ×*m* matrix, where *m* is 33,674 single nucleotide polymorphism (SNPs) and *q* is the 357 genotyped rice lines. Variance components and gBLUPs were obtained using ASREML (Release 4.0) (Gilmour et al., 2015).

Solving the mixed model equation will give three RR coefficients for each line. Using these RR coefficients, GEBVs at each time point can be obtained. For line *j* the predicted genetic values (GEBV) at each time point is given by *GEBV*_*j*_ = **Φ**_*g*_*û*_*j*_ (Mrode, 2014). **Φ**_*g*_ is the matrix of Legendre polynomials used for fitting the additive genetic effects. A detailed explanation of the RR model is provided in the appendix.

#### 2.2.2 Single time point

The following mixed model approach was used to fit gBLUPs at each time point

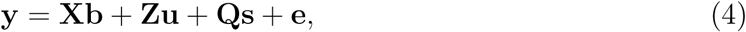

The matrices **X**, **Z** and **Q** correspond to incidence matrices for the fixed, random additive genetic and random experimental effect, respectively. Moreover, the dimensions for **X**, **Z** and **Q** are *n* × 1, *n* × *q* and *n* × *e*. We assume the random terms are distributed as follows 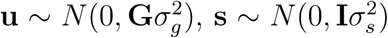, and 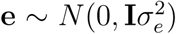. A genomic relationship matrix (**G**) was calculated as above and used for prediction of breeding values at each time point.

### 2.3 Genome-wide association analyses

#### 2.3.1 Estimating marker effects from GEBVs

GEBVs (**û**) can be parameterized as 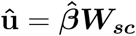, where **W_sc_** is a matrix of marker genotypes, as defined above, and 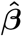 is a vector of allele substitution effects. 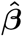 can be obtained using BLUP

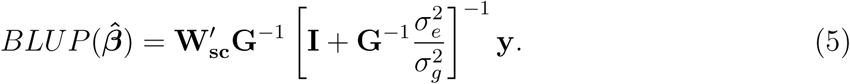

where 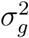 and 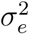 are genetic and residual variances, respectively.

Given BLUP of GEBVs is

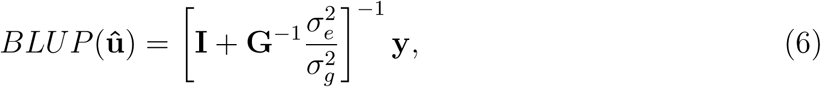

BLUP of marker effects can be obtained using the following linear transformation

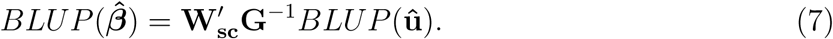

This relationship was leveraged to solve for marker effects from breeding values at each time point for both RR and single time point (TP) analyses.

#### 2.3.2 Variance of SNP effects

The variance of marker effects was calculated following the methods outlined by Duarte et al. (2014). Briefly, the variance of marker effects can be obtained via linear transformation of the variance of GEBVs (**û**).

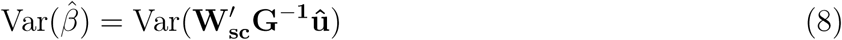

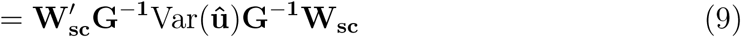

The prediction error variance (PEV) of **û** is

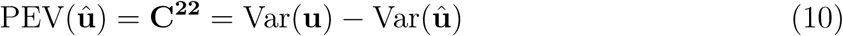

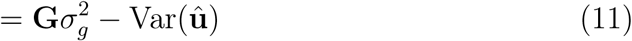

**C**^**22**^ is obtained by inverting the coefficient matrix of the mixed model equation provided in the appendix, and extracting the elements corresponding to additive genetic effects Henderson (1984). Thus, by rearranging equation 10, the variance of predicted breeding values is

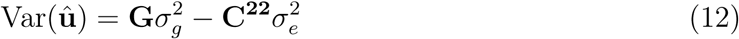

For the TP approach **C**^**22**^ is a *q* × *q* matrix, and diagonal elements correspond to the PEV of breeding values. Since the MME is solved for each time point independently, the above procedure can be used to obtain the variance of SNP effects on each day. However for the RR approach, **C**^**22**^ is *q* * *k*_*g*_ × *q* * *k*_*g*_ and represents the PEV for the additive genetic RR coefficients. Thus, to obtain Var(**û**) at each time point, we define a new matrix **C**^**22**^**\*** that is *q* * *d* × *q* * *d* where *d* is the number of time points (e.g. 20). This is given by

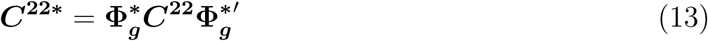

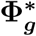 is a *q* * *d* × *k*_*g*_ * *q* block matrix where the diagonal sub-matrices consist of Legendre polynomials at each standardized time interval. This approach is analogous to that described by Mrode (2014), and is described in greater detail in the appendix.

#### 2.3.3 Obtaining p-values for marker effects

SNP effects for SNP_*j*_ at time *t* were divided by their corresponding 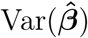 using

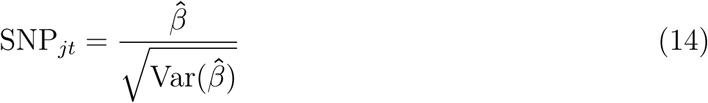

The *p*-values for marker effects were calculated as 1 minus the cumulative probability density of the absolute value of SNP_*jt*_, and this number was subsequently multiplied by two. This is summarized as follows.

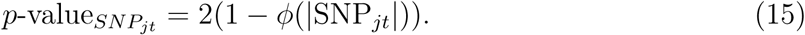

Following Zhao et al. (2011) a threshold of 1 × 10^−4^ was used to declare significant loci.

## 3 Results and Discussion

To identify loci associated with shoot growth trajectories in rice, we utilized a novel RR approach that allows for trait trajectories to be modeled across time points. Shoot growth trajectories were recorded for 357 diverse rice accessions over a period of 20 days during early vegetative growth (13 – 33 DAT). A RR model was fitted to the shoot growth trajectories, which included a fixed second-order Legendre polynomial, a random second-order Legendre polynomial for the additive genetic effect, a first-order Legendre polynomial for the environmental effect, and heterogeneous residual variances. GEBVs were predicted for each accession at each of the 20-time points as described in Campbell et al. (2018), and was used to estimate marker effects at each time point. Results from the RR were compared with a conventional single time point approach in which GEBVs were predicted at each time point using a conventional mixed model and were used to estimate marker effects.

### 3.1 RR-GWAS recovers more significant associations and increases predicted marker effect sizes

With RR models, the incorporation of the covariance structure of multiple measurements should lead to a more accurate partitioning of phenotypic variation into genetic and environmental components, and improve genetic inference. To demonstrate the advantages of a longitudinal genetic inference approach over a conventional TP approach, significant marker effects were compared between the RR and TP approaches. A 131% increase in the number of significant associations (*p* < 10^−4^) was observed with the RR approach compared to the conventional TP model. A total of 442 SNPs were found to be significantly associated with shoot growth trajectories at one or more time points using the RR approach, while 191 were found using the TP approach. Correlations in SNP effects estimated using the two approaches showed a very high agreement (*r* = 0.85), however predicted marker effects 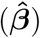 obtained using the RR were considerably larger than the single time point analysis (Fig 1). For instance, 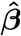 for the RR approach ranged from −299.1 to 295.0 across all days, while for the TP approach 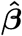 ranged from −104.6 to 112.3. These differences are evident in the distribution of marker effects pictured in Fig 1. Manhattan plots for each of the 20-time points is provided as supplemental Figures S2, S3, S6, S7, and the corresponding Q-Q plots are provided as supplemental Figures S10, S8, S9, S5. These results indicate that the utilization of information across all time points with the RR improves the ability to detect significant associations as well as increases the predicted marker effect sizes compared to a model that utilizes information at only a single time point.

**Figure 1:**
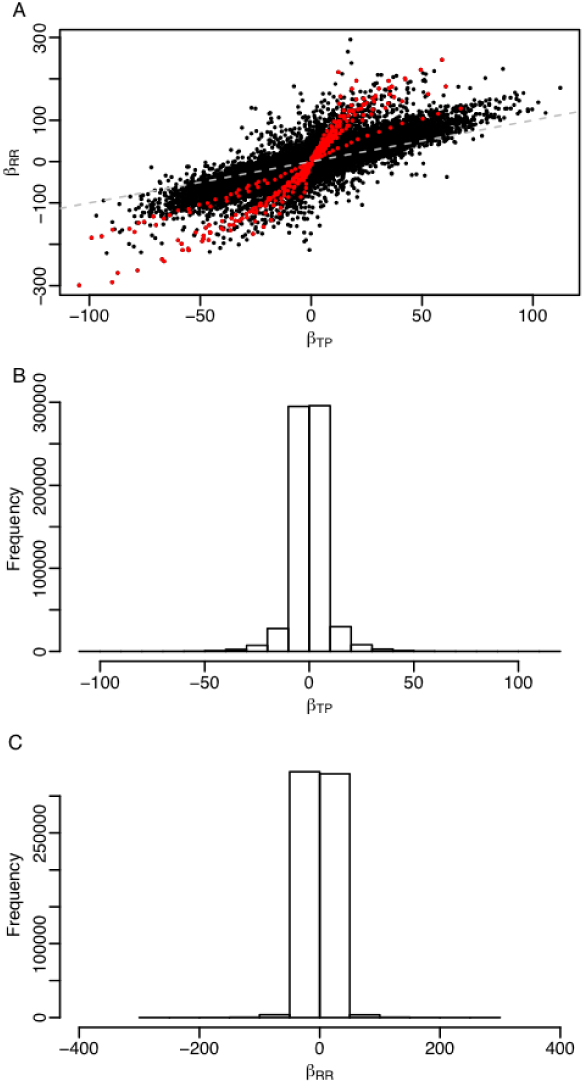
Correlation and distribution of SNP effects from random regression (RR) and single time point (TP) analysis. (A) Correlation between SNP effects for the random regression (*β*_*RR*_) and single time point analyses (*β*_*TP*_). SNPs highlighted in red are those that were statistically significant in the RR approach (*p* < 1 × 10^−4^). The grey broken lines depicts a one-to-one relationship between *β*_*RR*_ and *β*_*TP*_. Distribution of SNP effects across all 20-time points from the TP analyses (B) and RR analysis (C).

**Figure 2:**
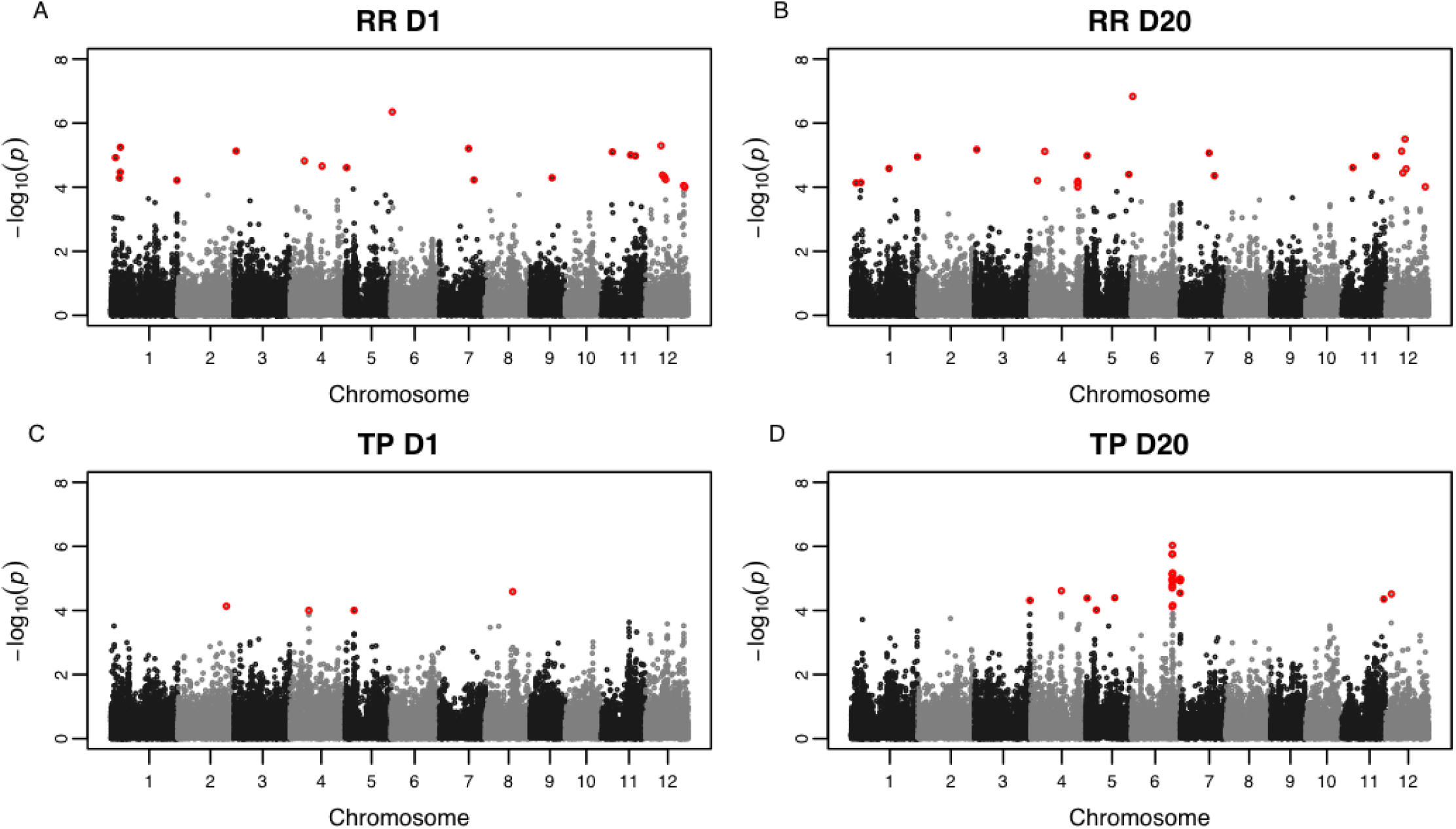
Manhattan plots for RR and TP approaches on days 1 and 20. (A,B) Manhattan plots for RR approach on days 1 and 20, respectively. (C,D) Manhattan plots for TP approach on days 1 and 20, respectively. −log_10_(*p*) is shown on the *y*-axis. Statistically significant SNPs are highlighted in red (*p* < 1 × 10^−4^).

These results suggest that the inclusion of time axis for genetic inference improve the ability to recover significant associations. Several other studies have showed similar improvements in the estimation of variance components and genetic inference using different approaches for longitudinal traits. For instance, De Andrade et al. (2002) showed a longitudinal approach that leveraged pedigree data and systolic blood pressure measurements collected at three time points improved heritability estimates compared with a single time point approach. While in the context of GWAS, Das et al. (2011) used a novel functional GWAS (*f* GWAS) approach and identified several new variants associated with body mass index collected at four time points in humans. Moreover, using simulated data the authors show that the statistical power exceeds 0.8 with a false positive rate of less than 0.1 for sample sizes greater than 1,000. Similar gains for GWAS have been demonstrated in plants, animals, and humans (Xu et al., 2014; Campbell et al., 2015; Yi et al., 2015; Lund et al., 2008).

### 3.2 RR-GWAS reveals the dynamic genetic architecture of shoot growth responses in rice

For many traits, such as growth, genetic effects are expected to vary across time. These temporal genetic effects can be effectively captured using a RR approach. To examine the dynamic genetic architecture of shoot growth trajectories, significant SNPs from the RR approach were selected and those within a 200 kb window were merged to a single QTL. The 200 kb window that we used corresponds to the average linkage disequilibrium in rice (Zhao et al., 2011; Huang et al., 2010). For the RR approach, a total of 26 significant QTL were detected at one or more time points, while for the TP approach only 15 significant QTL were detected.

To dissect the dynamic genetic architecture of shoot growth in rice, significant QTLs were classified into four categories: persistent QTL (QTL detected at all 20-time points), long-duration QTL (those with significant associations at more than 12, but less than 20-time points), mid-duration (QTL with associations at 6 – 12 time points), and short-duration QTL (those with associations at fewer than 6-time points). Of these categories, far more persistent QTLs were detected, with a total of 13 observed at all 20-time points. Short duration QTL also showed the fewest number of significant QTL (2). While five and six QTL were detected for long and mid-duration QTL, respectively. The frequencies of significant QTL for each category were calculated at each time point and plotted as a function of time (Fig S7). The majority of long-duration QTL were detected towards the end of the experiment (day 8 onward), while short-duration QTL were detected only from days 1 to 4. Mid-duration QTL were detected at all time points. The *p*-values across all 20-time points for all significant QTL are provided in Figure 3. Collectively, these results indicate that the shoot growth is regulated by numerous loci that have both transient and persistent effects throughout early vegetative growth.

**Figure 3:**
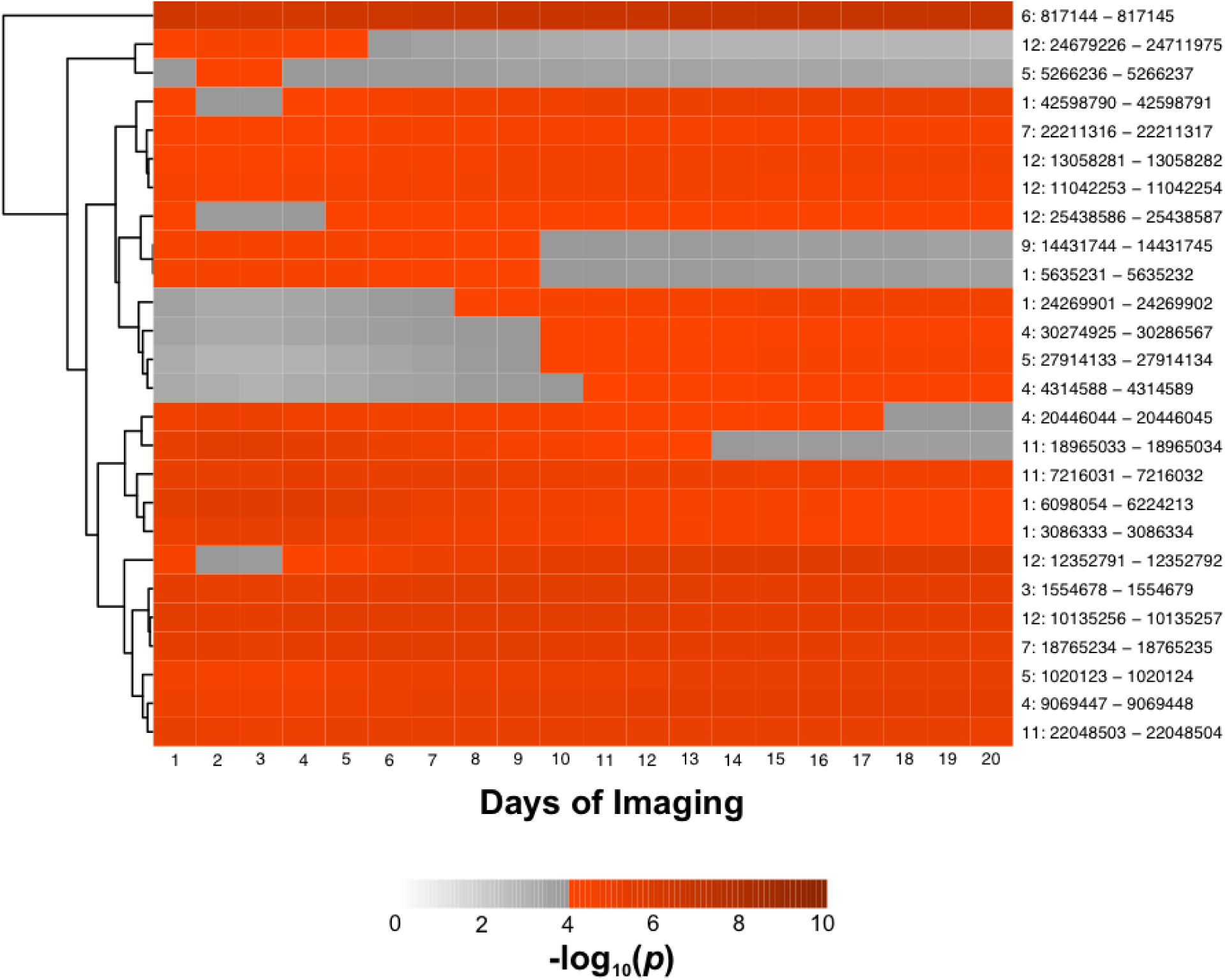
Heatmap showing time-specific QTL. A subset of significant QTL identified with RR approach are pictured. The *x*-axis indicates the days of imaging and the *y*-axis shows the chromosome and intervals for the QTL. For each QTL, the most significant SNP within the interval at each time point were selected. The grey color scale indicates a non-significant association, while the red color scale indicates a statistically significant association (*p* < 1 × 10^−4^).

The importance of time-specific QTL has been demonstrated in both plants and animals (Moore et al., 2013; Bac-Molenaar et al., 2015; Campbell et al., 2015, 2017). For instance, using a single time point linkage mapping approach, Moore et al. (2013) showed several time-specific QTL associated with root gravitropic responses in Arabidopsis. Moreover, many of these QTL harbored candidate genes known to influence root growth, root gravitropism, or hormone transport and signaling. Bac-Molenaar et al. (2015) collected rosette growth trajectories over a period of 20 days for a diverse panel of 324 Arabidopsis accessions. A growth function was fit for each accession, and model parameters were used for GWAS. The authors showed that many associations detected for model parameters were also detected at a few time points using a single time point GWAS approach. While few longitudinal studies have been performed in rice, our previous studies have identified time-specific QTL for shoot growth and salt stress responses (Campbell et al., 2015, 2017).

## 4 Conclusion

New phenotyping platforms have provided the plant science community with a suite of tools to collect high-dimensional temporal phenotypic data. With these temporal dataset, quantitative genetic approaches that can leverage the covariance across time points must be fully utilized to realize the potential of these data for genomic prediction and genetic inference. In this study, we show that the RR framework that has been extensively developed in animal breeding can be extended to genetic inference in plants. This approach can effectively be used to identify QTL with time-specific effects. To date, this is the first application of random regression models for genetic inference of a longitudinal trait in a major crop.

## Acknowledgements

Funding for this research was provided by the National Science Foundation (United States) through Award No. 1238125 to Harkamal Walia, and Award No. 1736192 to Harkamal Walia and Gota Morota.

## Supplemental Materials

Supplemental Data.pdf: Appendix; Figures S1-S6.

## Author Contributions

Study was conceived by H.W., G.M., and M.C.; phenotyping was performed by M.C. and H.W.; M.C., M.M. and G.M. performed all analyses; M.C. wrote the manuscript, and editorial comments were provided by M.M., H.W. and G.M.

## Data Accessibility

The full datasets and all code used in this study is available via GitHub (https://github.com/malachycampb RR-GEBVs-for-genomic-inference-of-longitudinal-traits) and the WRCHR website (WRCHR.org).

## Supplemental Data

**Figure S1:**
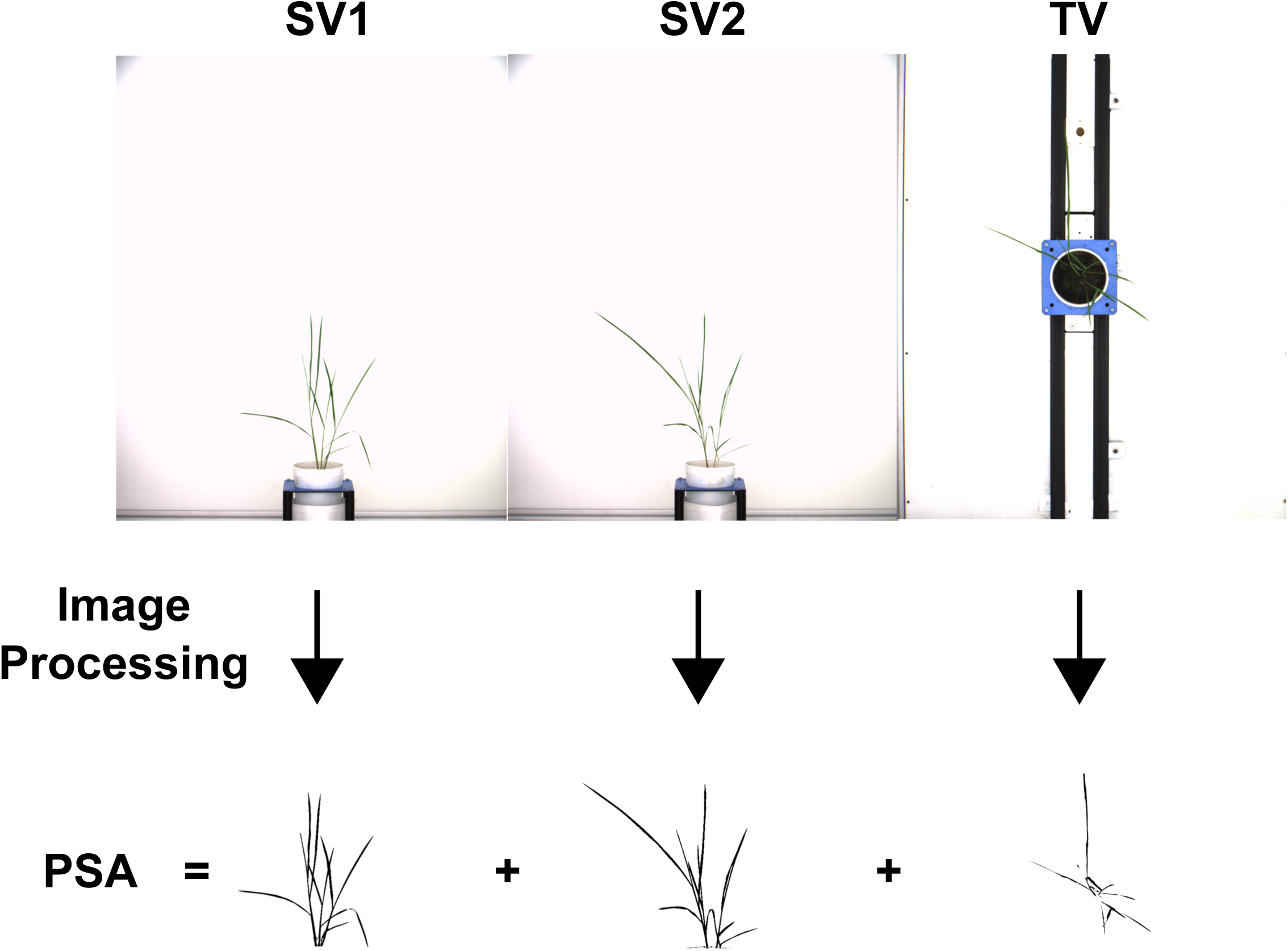
Visual depiction of assessing shoot biomass using projected shoot area. The plant is imaged from three perspectives. The two side view (SV) images are separated by 90 degrees. TV: top view; PSA: projected shoot area

**Figure S2:**
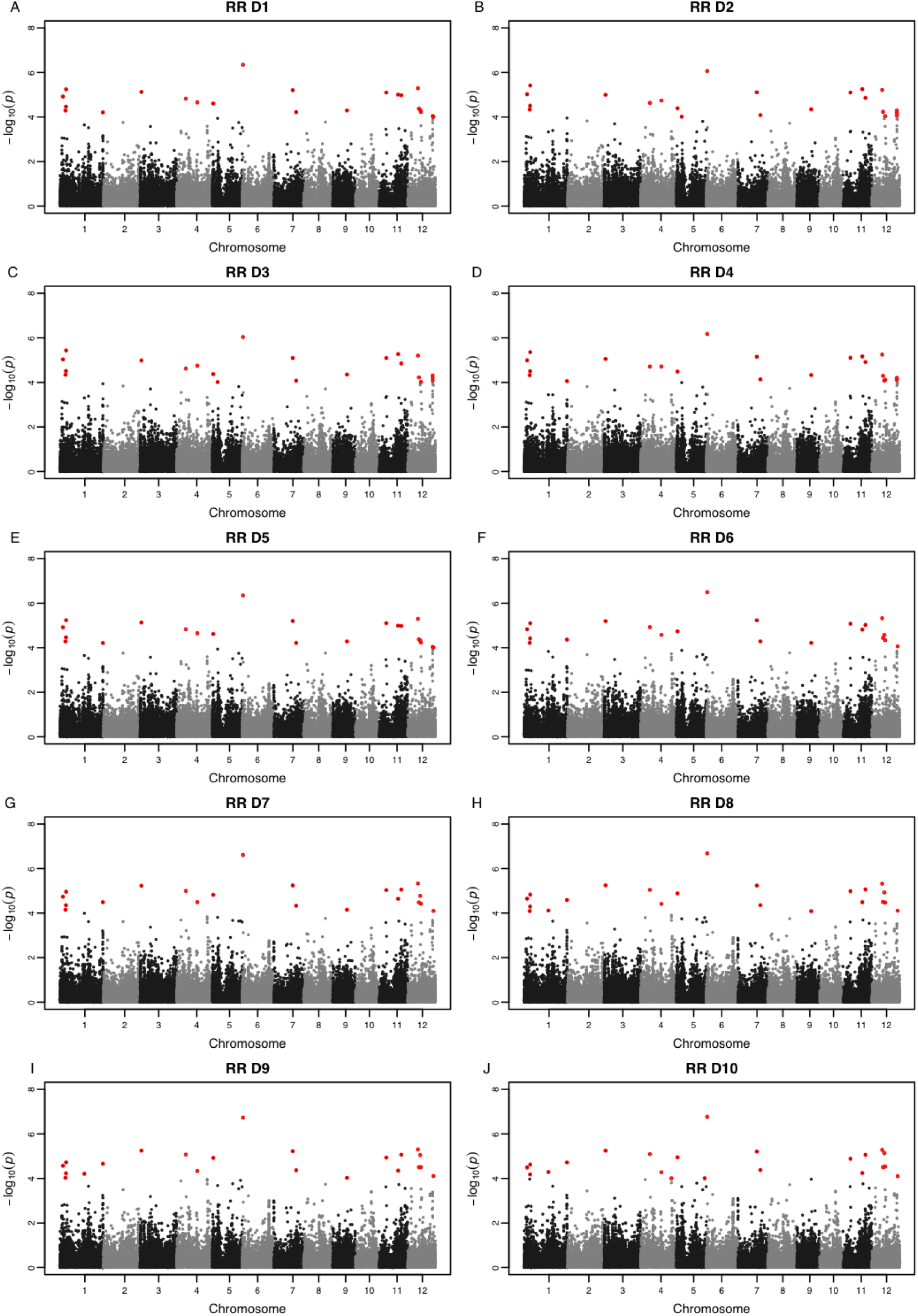
Manhattan plots for RR approach at days 1 to 10. Each panel represents a single time point. −log_10_(*p*) is shown on the *y*-axis. Statistically significant SNPs are highlighted in red (*p* < 1 × 10^−4^).

**Figure S3:**
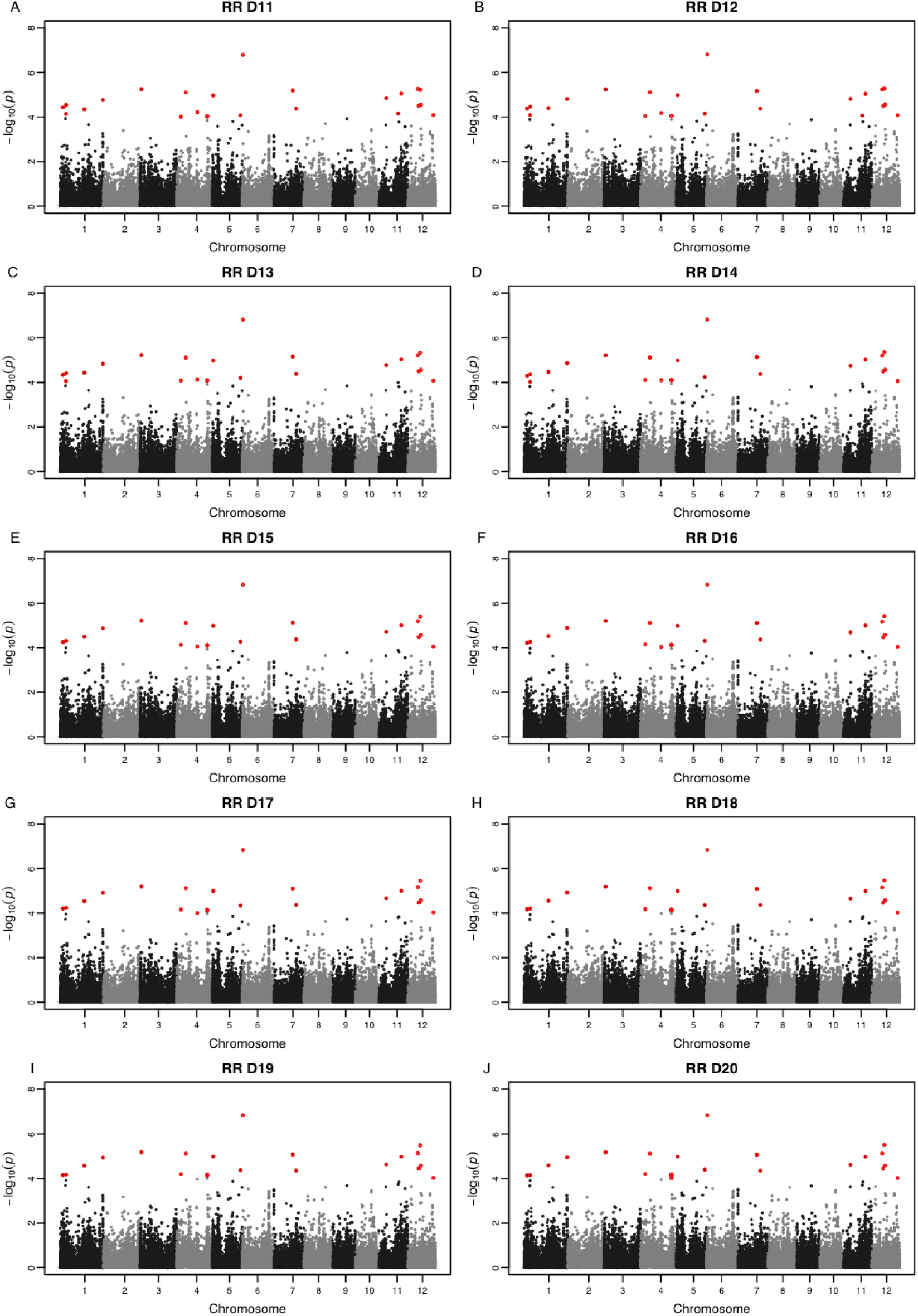
Manhattan plots for RR approach at days 10 to 20. Each panel represents a single time point. −log_10_(*p*) is shown on the *y*-axis. Statistically significant SNPs are highlighted in red (*p* < 1 × 10^−4^).

**Figure S4:**
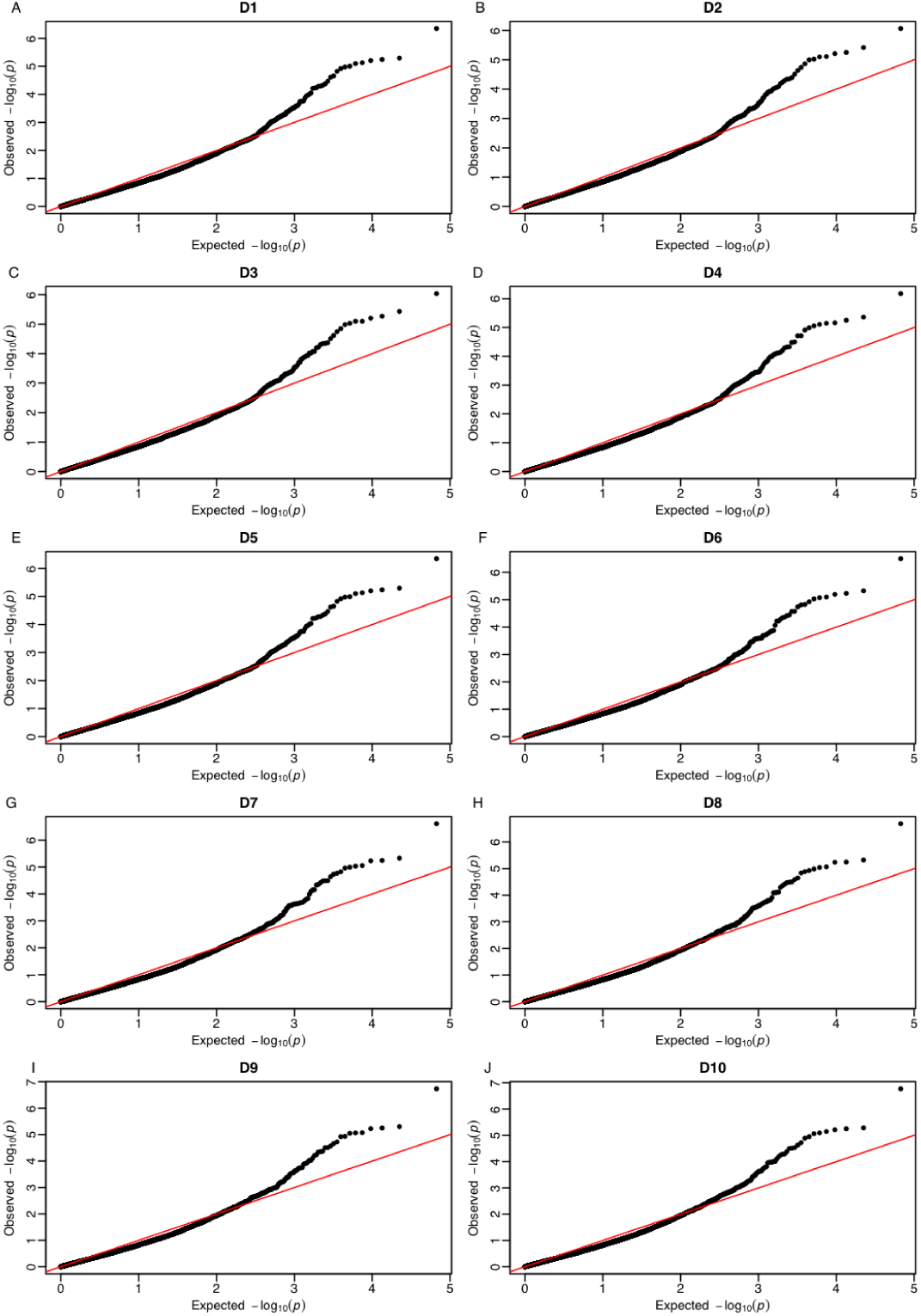
QQ plots for RR approach at days 1 to 10. Each panel represents a single time point. The observed −log_10_(*p*) is shown on the *y*-axis, while the expected −log_10_(*p*) is shown on the x-axis.

**Figure S5:**
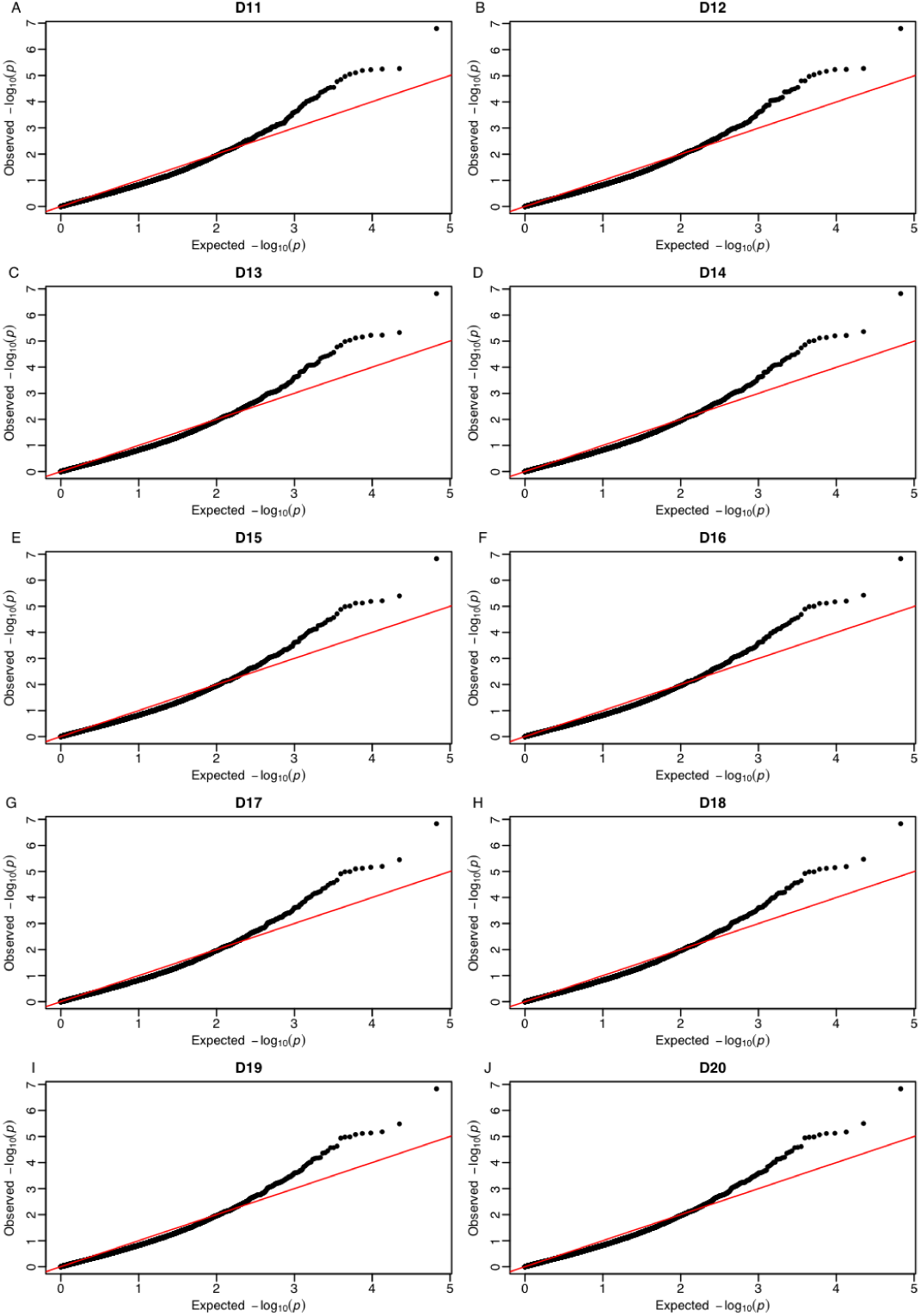
QQ plots for RR approach at days 11 to 20. Each panel represents a single time point. The observed −log_10_(*p*) is shown on the *y*-axis, while the expected −log_10_(*p*) is shown on the x-axis.

**Figure S6:**
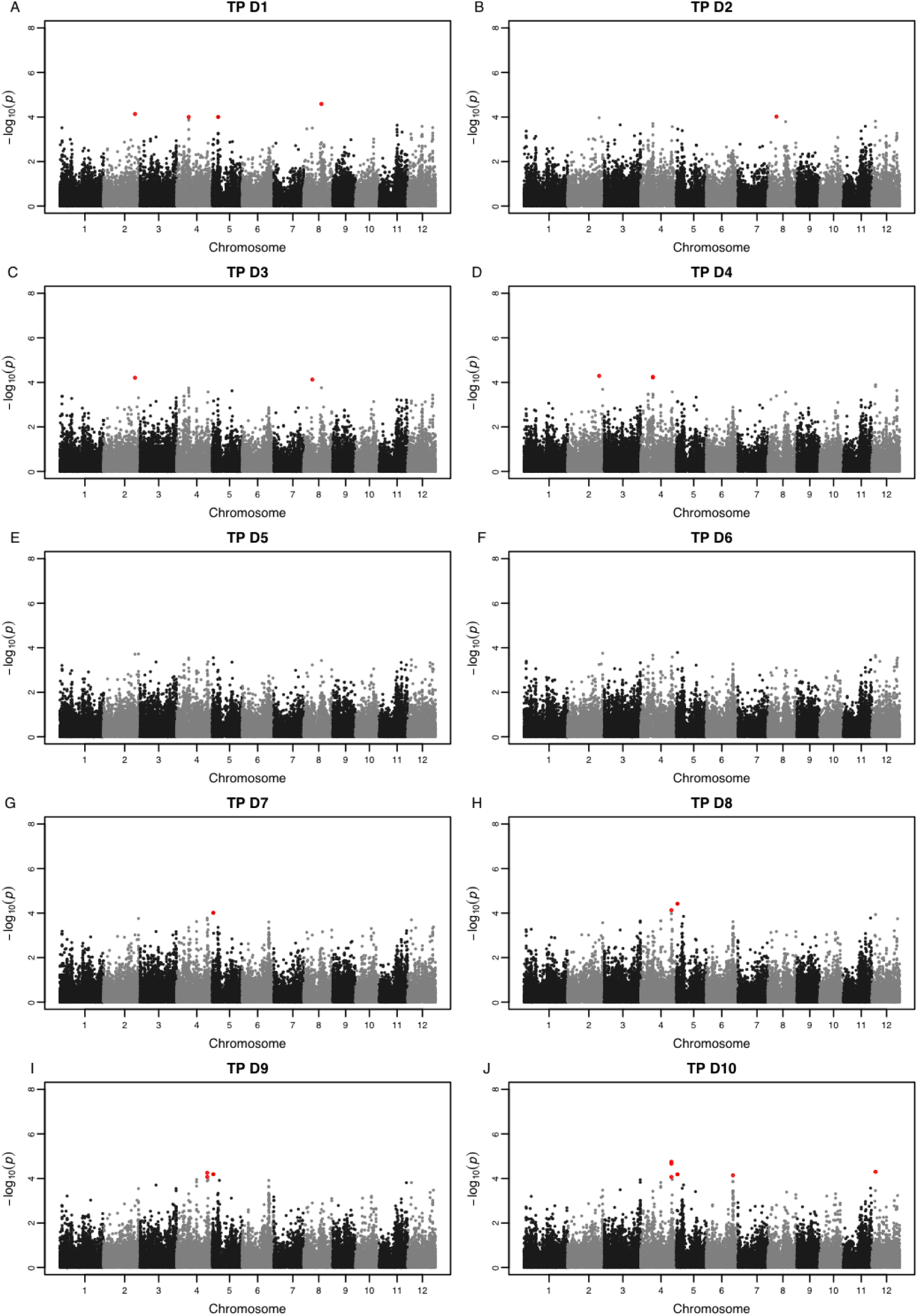
Manhattan plots for TP approach at days 1 to 10. Each panel represents a single time point. −log_10_(*p*) is shown on the *y*-axis. Statistically significant SNPs are highlighted in red (*p* < 1 × 10^−4^).

**Figure S7:**
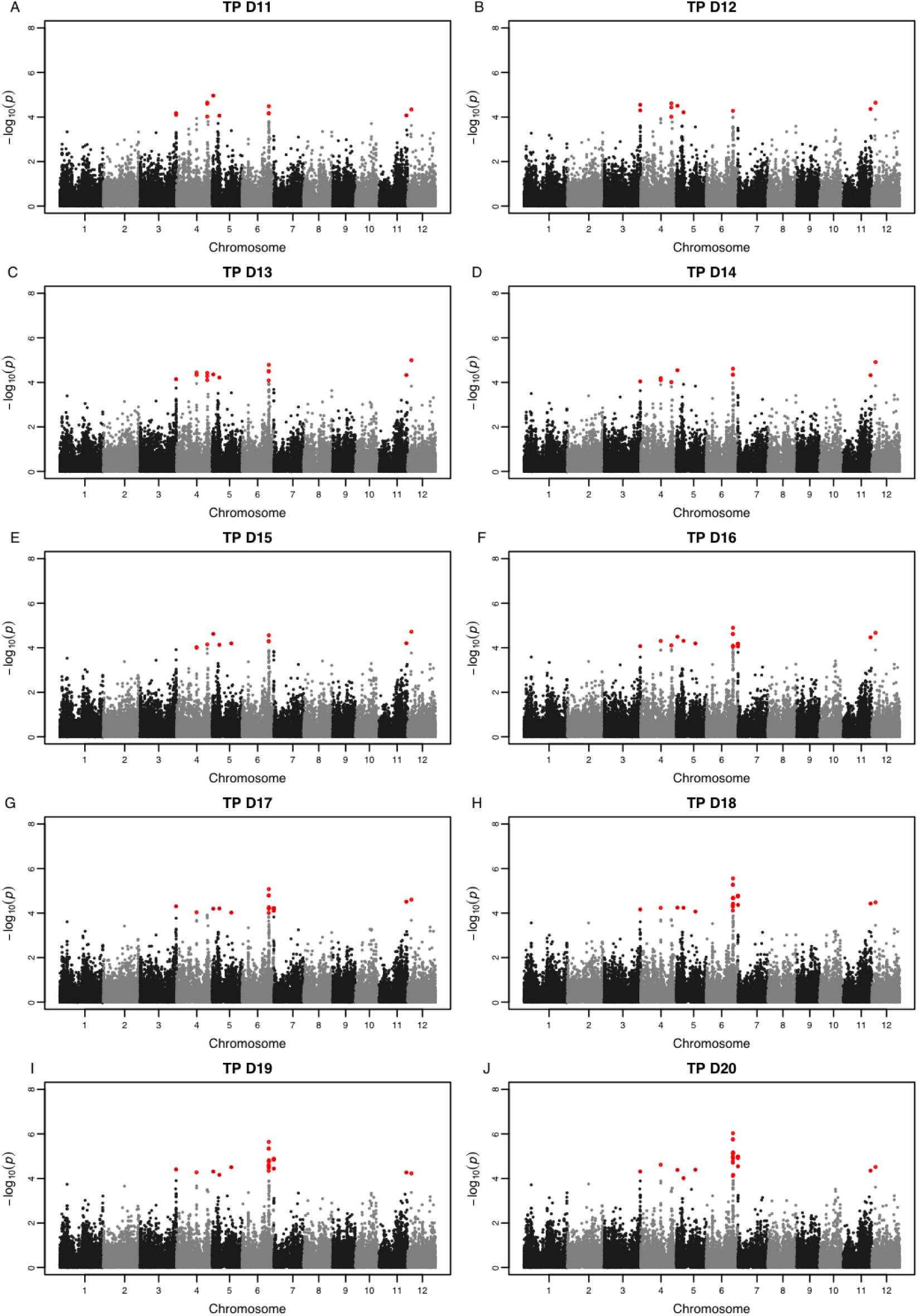
Manhattan plots for TP approach at days 11 to 20. Each panel represents a single time point. −log_10_(*p*) is shown on the *y*-axis. Statistically significant SNPs are highlighted in red (*p* < 1 × 10^−4^).

**Figure S8:**
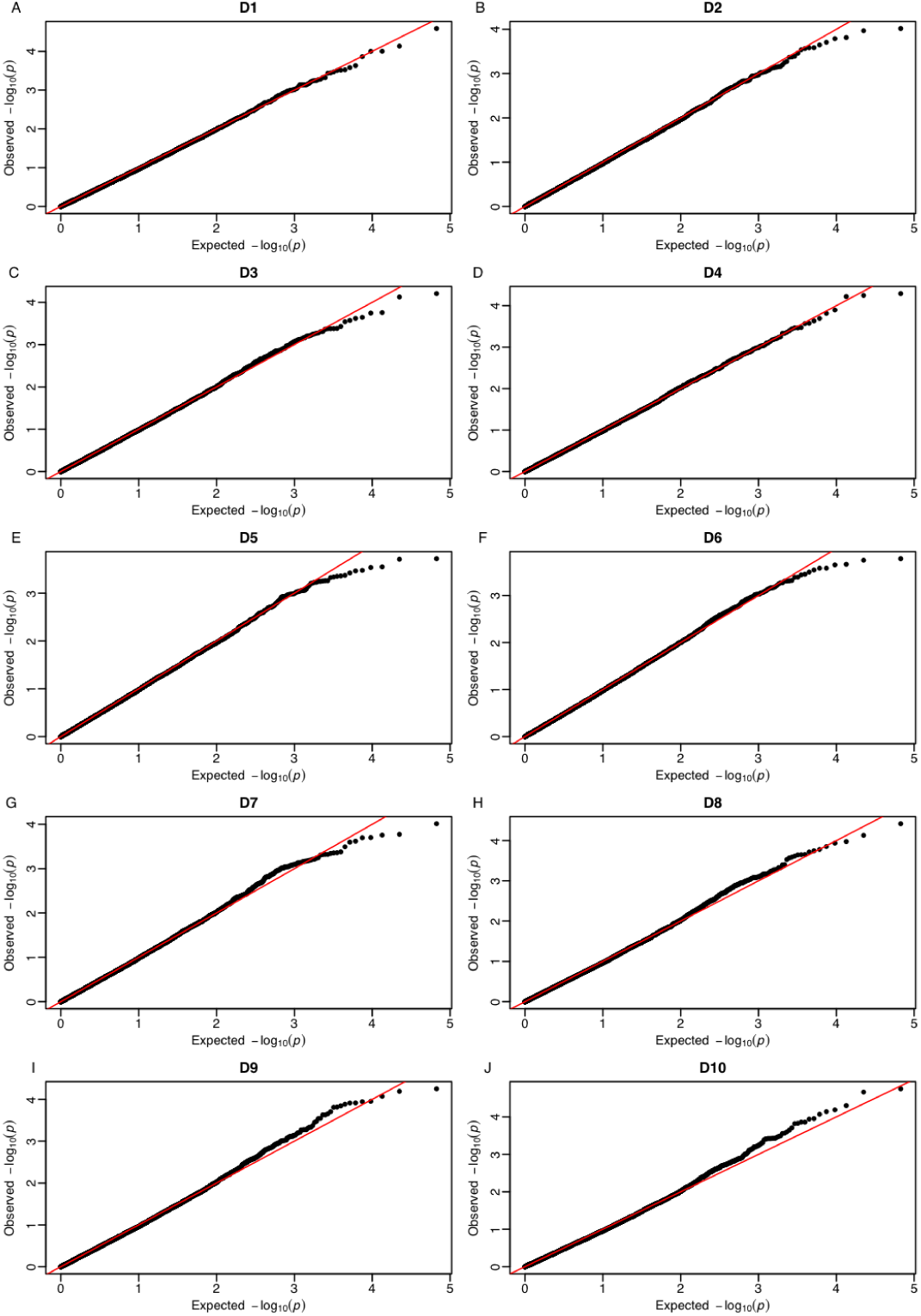
QQ plots for TP approach at days 1 to 10. Each panel represents a single time point. The observed −log_10_(*p*) is shown on the *y*-axis, while the expected −log_10_(*p*) is shown on the x-axis.

**Figure S9:**
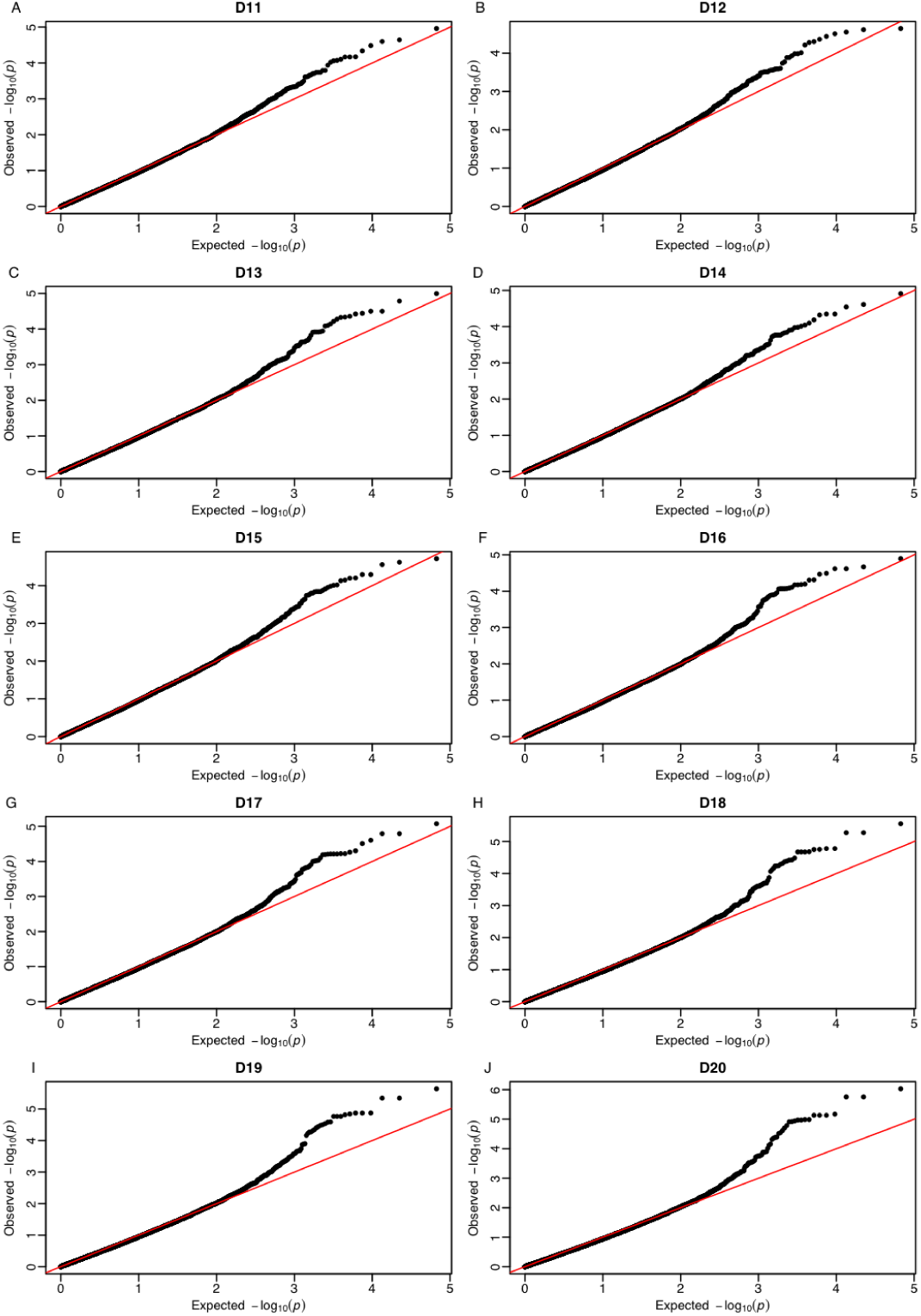
QQ plots for TP approach at days 11 to 20. Each panel represents a single time point. The observed −log_10_(*p*) is shown on the *y*-axis, while the expected −log_10_(*p*) is shown on the x-axis.

**Figure S10:**
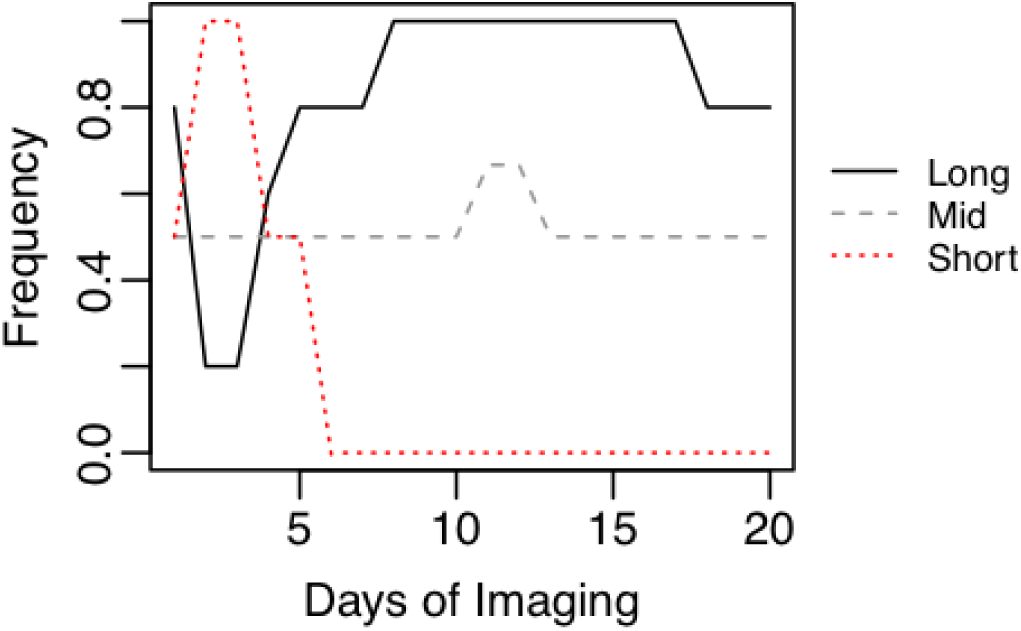
Frequency of time-specific QTL. Long refers to long-duration QTL that were detected on more than 12, but less than 20 days. Mid refers to mid-duration QTL that were detected between 6 to 12-time points. Short indicates short-duration QTL, which were detected at fewer than 6-time points. Frequency was determined by dividing the number of QTL detected at time *t* by the total number of QTL for a given class.

## Appendix

### Random regression gBLUP using Legendre polynomials

Polynomial functions are an attractive approach to model longitudinal data, as they require no prior knowledge of the shape of trait trajectories and can be estimated using linear modeling approaches. However, there is often a high correlation between components of the polynomial function. Orthogonal polynomials on the other hand, such as Legendre polynomials, have the same attractive characteristics of polynomial functions and also reduce the correlation between polynomial components. Legendre polynomials are defined on a standardized time interval [−1, 1] using 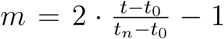, where *t*_0_ is the first time point with data, and *t*_*n*_ is the last time point in the data set (Kirkpatrick et al., 1990, 1994).

Consider a simple case where we wish to partition a process measured over three time points (*y*) into fixed (*µ*) and random (*α*) time dependant effects. The RR model can be defined as *y*(*t*) = *µ*(*t*) + *α*(*t*) + *ϵ*. We can obtain a “full” fit using a second-order Legendre polynomial. The first two Legendre polynomials are *P*_*o*_(*x*) = 1 and *P*_1_(*x*) = *x*. Subsequent polynomials can be calculated using 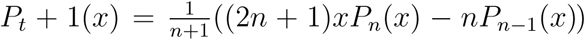. Thus, for *P*_2_(*x*) the Legendre polynomial is 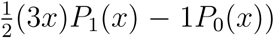. These Legendre polynomials are then normalized using 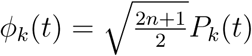, giving *ϕ*_0_(*t*) = 0.7071, *ϕ*_1_(*t*) = 1.2247(*t*) and *ϕ*_2_(*t*) = 2.317(*t*^2^) – 0.7906. Two matrices can be defined, **Λ** and **M**, that store the coefficients for the Legendre polynomials and the standardized time values, respectively.

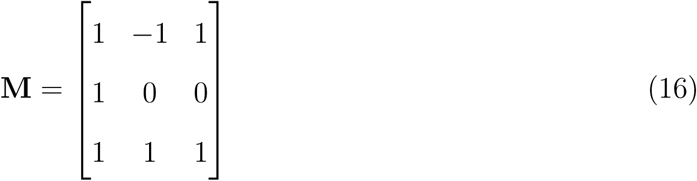

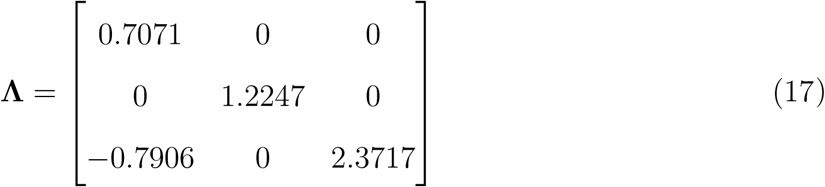

Multiplying the two gives **Φ** where each row vector corresponds to the series of Legendre polynomials at each standardized time interval.

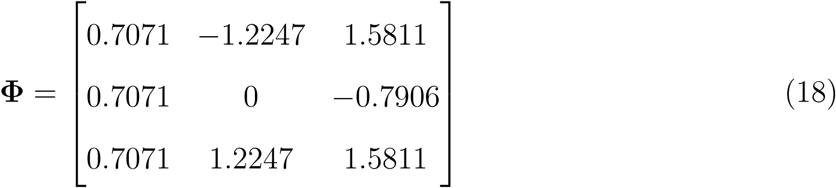

The covariance matrix for the RR coefficients is given by **K**. The full covariance matrix (**V**) among all three time points can be obtained via **V** = **ΦKΦ′**.

In the following study we aimed to assess the genetic and environmental covariances for shoot growth measured across a period of 20-time points. To this end, we utilized a RR model that modeled the fixed population mean (*β*) growth trajectories using a secondorder Legendre polynomial, and the random genetic (*u*) and experimental effects (*s*) using a second-order and first-order Legendre polynomial respectively. Following the example above, these time-dependant processes can be described using a linear combination of **Φ**. The covariances at each time point for the random genetic and experimental effects are given by **V**_*g*_ = **Φ**_*g*_**ΩΦ*g*′** and **V**_*s*_ = **Φ**_*s*_**PΦ**_*s*_**′**, respectively. The matrices **Ω** and **P** represent the covariance matrices for the RR coefficients for the genetic and experimental effects, respectively. Thus, the dimensions of **Ω** 3 × 3 and **P** is 2 × 2.

#### Defining the mixed model equation

The following random regression model was used to model trajectories for PSA across the 20-time points and obtain estimates for **Ω** and **P**

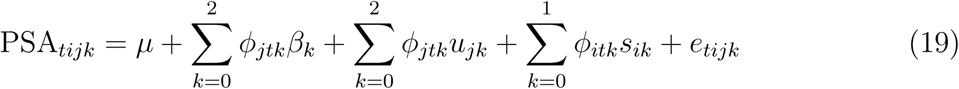

In matrix notation, the model can be written as

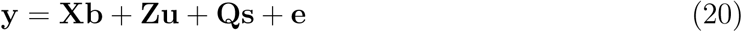

**y** is a vector with an order equal to the number of observations and contains the *PSA* over the 20 days. **X** is an covariable matrix for the fixed effects where the number of rows is equal to the number of observations (*n*) and the number of columns is equal to the order of Legendre polynomial used to model fixed effects (*k*_*f*_). The matrices **Z** and **Q** are covariable matrices for the random additive genetic and random experimental effects, respectively. The number of rows for **Z** is equal to the number of observations and the number of columns corresponds to the order of Legendre polynomial times the number of lines used to fit the additive genetic effect (*q* * *k*_*g*_ = 357 * 3 = 1, 071). For **Q** the number of columns would be 6 (*e* * *k*_*s*_ = 3 * 2) and the number of rows would be equal to the number of observations. We assume **u** ∼ *N* (0, **G** ⊗ **Ω**), **s** ∼ *N* (0, **I** ⊗ **P**), and **e** ∼ *N* (0, **I** ⊗ **D**). Here, **Ω** and **P** are the covariance matrices for the RR coefficients for the additive genetic and permanent environmental effects. **D** is a diagonal matrix that allows for heterogeneous variances over the 20-time points.

The mixed model equation (MME) is

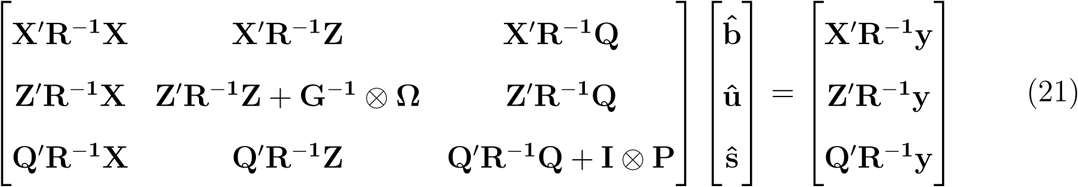

Solving the above MME will give three RR coefficients for each line for the random genetic effects. Using these RR coefficients, the genetic values at each time point can be obtained as described above. For line *j*, the predicted genetic values (gBLUP) at each time point is given by *gBLUP*_*j*_ = **Φ***û*_*j*_.

### Constructing the covariable matrices

For each term, we define a matrix of Legendre polynomials evaluated at each time point. Recall that both the fixed and random additive genetic effect are modeled using a second-order Legendre polynomial. Thus the matrix of Legendre polynomials for the fixed and random additive genetic effect for the first and last three time points is, (**Φ*f***, Φ***g***, respectively) are

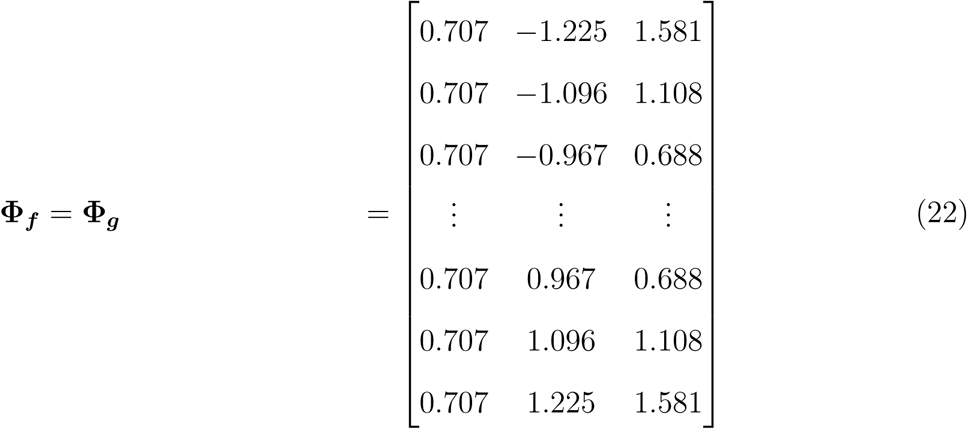

For the environmental effect, the matrix of Legendre polynomials (**Φ**_*s*_) is of order *t* × 2 and for the first and last three time points is

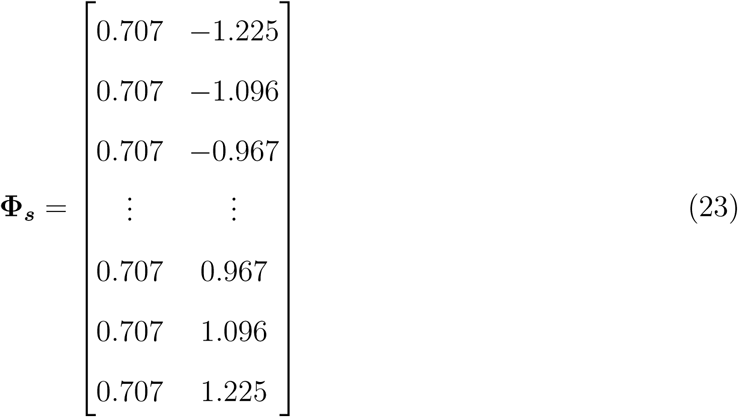

The covariable matrix **X** is defined as **X** = **X°Φ_f_** where **X°** is a vector of 1 with length *q* * *e*. Similarly, we define matrices **Z** and **Q** as

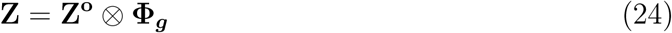

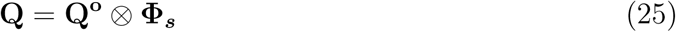

**Z°** and **Q°** are incidence matrices that allocate temporal records to individuals and experiments respectively. The order of **Z°** is *q* * *e* × *q* and **Q°** is *q* * *e* × *e* (*q* is the number of individuals, *e* is the number of experiments).

### Calculating Var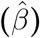 at each time point

The objective is to calculate SNP effects at each time point. Recall for a univariate gBLUP approach (e.g. the single time point approach), SNP effects can be obtained from breeding values through a simple linear transformation given by

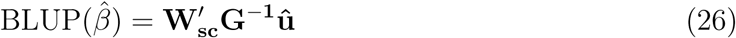

Thus,

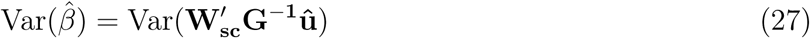

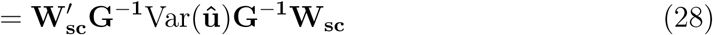

The prediction error variance (PEV) of ***û*** is

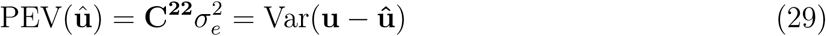

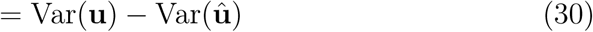

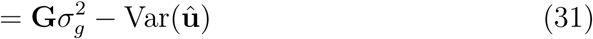

By rearranging the equation above we obtain

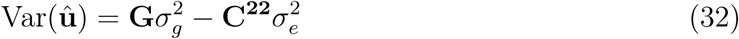

To calculate the variance of the SNP effects, we can introduce equation 34 into equation 27 giving

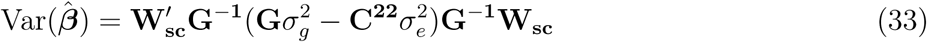

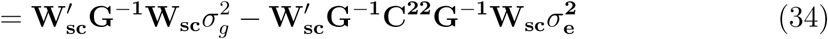

At each time point, 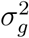 is extracted from the corresponding the diagonal element of the matrix of genetic covariances for each time point, given by 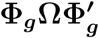.

**C**^**22**^ is obtained by inverting the coefficient matrix of the MME (equation 21), and is of order *q* * *k*_*g*_ × *q* * *k*_*g*_. The diagonal elements of **C**^**22**^ contain the PEV for the RR coefficients for the additive genetic effect. To obtain the variance of SNP effects at each time point, **C**^**22**^ must be transformed so that the diagonal elements correspond to the PEV for GEBVs at each time point. We will refer to this *q* * *d* × *q* * *d* matrix as **C**^**22**^**\***. Following (Mrode, 2014), PEV for individual *i* at each time point can be obtained by taking the diagonal elements of 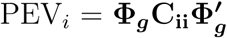. **C**_**ii**_ is a 3 × 3 submatrix of RR coefficients from **C**_**22**_ for individual *i*. To extend this approach to the full **C**_**22**_ matrix, we construct a block matrix of **Φ*g*** 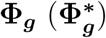 via 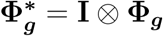 and obtain ***C*^22^\*** by

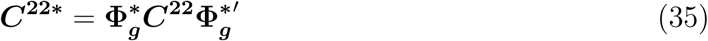

Thus, 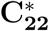 is *q* * *t* × *q* * *t* and the diagonal elements are the PEV for GEBVs at each time point. Finally, to calculate the variance of SNP effects at each time point at each, we extract the corresponding elements of ***C***^**22**^**\*** and introduce them into **??**. Calculation of *p*-values from this point is straight forward. ***C***^**22**^ for each time point can be extracted from ***C***^**22**^**\***, and calculation of *p*-values follows the procedures outlined in Materials and Methods.

## Abbreviations

BLUP: best-linear unbiased prediction
GEBVs: Genomic estimated breeding values
GWAS: genome-wide association study
PSA: projected shoot area
QTL: quantitative trait loci
RDP1: rice diversity panel 1
DAT: days after transplant
RR: random regression
SMR: single marker regression
SNP: single nucleotide polymorphism
TP: single time point

